# HSP90 interacts with VP37 to facilitate the cell-to-cell movement of broad bean wilt virus 2

**DOI:** 10.1101/2024.08.15.608091

**Authors:** Myung-Hwi Kim, Seok-Yeong Jang, Ji-Soo Choi, Sora Kim, Yubin Lee, Suejin Park, Sun-Jung Kwon, Jang-Kyun Seo

## Abstract

The systemic spread of viruses in plants requires successful viral cell-to-cell movement through plasmodesmata (PD). Viral movement proteins (MPs) interact with cellular proteins to modify and utilize host transport routes. Broad bean wilt virus 2 (BBWV2), a widespread plant RNA virus, moves from cell to cell as a virion through the PD gated by VP37, the MP of BBWV2. However, the host proteins that function in the cell-to-cell movement of BBWV2 remain unclear. In this study, we identified cellular heat shock protein 90 (HSP90) as an interacting partner of VP37. The interaction between HSP90 and VP37 was assessed using the yeast two-hybrid assay, co-immunoprecipitation, and bimolecular fluorescence complementation. Tobacco rattle virus-based virus-induced gene silencing analysis revealed that *HSP90* silencing significantly inhibited the systemic spread of BBWV2 in *N. benthamiana* plants. Furthermore, *in planta* treatment with geldanamycin (GDA), an inhibitor of the chaperone function of HSP90, demonstrated the necessity of HSP90 in successful cell-to-cell movement and systemic infection of BBWV2. Interestingly, GDA treatment inhibited the HSP90-VP37 interaction at the PD, resulting in the inhibition of VP37-derived tubule formation through the PD. Our results suggest that the HSP90-VP37 interaction regulates VP37-derived tubule formation through the PD, thereby facilitating the cell-to-cell movement of BBWV2.

**IMPORTANCE:** This study highlights the regulatory role of heat shock protein 90 (HSP90) in facilitating the cell-to-cell movement of broad bean wilt virus 2 (BBWV2). HSP90 interacted with VP37, the movement protein of BBWV2, specifically at plasmodesmata (PD). This study demonstrated that the HSP90-VP37 interaction is crucial for viral cell-to-cell movement and the formation of VP37-derived tubules, which are essential structures for virus transport through the PD. The ATP-dependent chaperone activity of HSP90 is integral to this interaction, as demonstrated by the inhibition of virus movement upon treatment with geldanamycin (GDA), which disrupts the function of HSP90. These findings elucidate the molecular mechanisms underlying the cell-to-cell movement of plant viruses, and highlight the role of HSP90 in viral infection. This study suggests that the chaperone activity of HSP90 may function in changing the conformational structure of VP37, thereby facilitating the assembly and function of virus-induced structures required for viral cell-to-cell movement.

## Introduction

Plant viruses are obligate intracellular parasites that depend on living host cells for proliferation. They induce varying degrees of biotic stress by hijacking host cellular machinery and disturbing metabolic functions during infection, resulting in physiological and morphological changes on host plants (1–3). In addition, infection involves multiple compatible interactions between viral and host proteins at various stages of the viral life cycle, including gene expression, replication, and spread (4–6). Thus, interactions between viral and host proteins have markedly influenced the evolution of plant viruses (2).

Viral transport within plants relies on non-structural proteins known as movement proteins (MPs). In most plant viruses, MPs orchestrate intracellular, intercellular, and long-distance movement to facilitate systemic infection (7). Although viral MPs are structurally diverse, they share common characteristics, such as localization to plasmodesmata (PD), PD gating, and interaction with host proteins during these processes (8, 9). The reliance of viral movement on host proteins is strongly associated with the physical interactions between viral MPs and host proteins. Therefore, identifying host proteins that interact with viral MPs is crucial for elucidating the molecular mechanisms underlying the intracellular and intercellular movements of plant viruses. Accumulating evidence reveals that viral MPs interact with various host proteins associated with PD and cellular transport systems to modify and utilize existing transport routes in plants (6, 10, 11).

Broad bean wilt virus 2 (BBWV2), a member of the genus *Fabavirus* in the family *Secoviridae*, is an emerging viral pathogen with a broad host range. It poses a significant threat to various economically important crops, including pepper and legume (12). The BBWV2 genome consists of two single-stranded positive-sense RNAs (RNA1 and RNA2) that are approximately 5,960 and 3,600 nucleotides long, respectively (13). RNA1 contains a single open reading frame that encodes one polyprotein precursor. This precursor is proteolytically cleaved to produce five mature proteins: the protease cofactor, NTP-binding motif, viral genome-linked protein, protease, and RNA-dependent RNA polymerase. RNA2 encodes two largely overlapping polyproteins using two alternative in-frame start codons. These polyproteins differ only at their N-termini and are proteolytically cleaved at the identical cleavage sites to produce four mature proteins: a replication-associated protein of 53 kDa (VP53), a viral movement protein of 37 kDa (VP37), as well as large (LCP) and small (SCP) coat proteins (13).

The role of VP37 in viral infection has been extensively demonstrated (13–16). VP37 is localized to the PD and forms tubule structures that facilitate the tubule-guided cell-to-cell movement of BBWV2 virions (13, 16). In addition, it can bind single-stranded nucleic acids and suppress RNA silencing (17). However, the host cellular proteins that interact with VP37 are yet to be identified. In our previous study, we engineered an infectious cDNA construct of BBWV2 RNA2 to express VP37 tagged with a Flag epitope (Asp-Tyr-Lys-Asp-Asp-Asp-Asp-Asp-Lys) at the C-terminus (VP37-Flag) during BBWV2 replication (13, 18). We also demonstrated that tagging VP37 with a Flag epitope at the C-terminus did not impair VP37 function in viral infectivity (13). Therefore, the present study aimed to apply this approach for an immunoprecipitation assay followed by liquid chromatography coupled with tandem mass spectrometry (LC-MS/MS) to identify the host cellular proteins that interact with VP37. Based on this approach and additional experiments, we identified cellular heat shock protein 90 (HSP90) as an interacting partner of VP37 in *Nicotiana benthamiana*. Our study also revealed that the HSP90-VP37 interaction occurred at the PD and was essential for the cell-to-cell movement of BBWV2, suggesting that HSP90 is a host factor that regulates BBWV2 movement through its interaction with VP37.

## Materials and Methods

### Plant growth and agroinfiltration

*N. benthamiana* plants were grown in an insect-free growth chamber under controlled conditions, with a 16-h light period at 26 °C followed by an 8-h dark period at 24 °C. For agroinfiltration, T-DNA-based binary vector constructs were transformed into *Agrobacterium* strain EHA105. Next, the *Agrobacterium* cultures carrying each construct were grown overnight at 30 °C in YEP medium supplemented with kanamycin (100 µg/mL) and acetosyringon (20 μM). The cultures were then centrifugated at 3000 rpm for 10 min, and the resulting bacterial pellet was resuspended to 0.5 OD_600_ in an infiltration buffer (MS salts, 10 mM MES, pH 5.6, 200 μM acetosyringon). The resuspended Agrobacteria were incubated at 30 °C for 4 h and infiltrated into the abaxial surface of *N. benthamiana* leaves using a 1-mL syringe. Each inoculation experiment was repeated independently at least thrice, with at least three plants per construct in each experiment.

### Immunoprecipitation and liquid chromatography coupled with tandem mass spectrometry (LC-MS/MS) analysis

*N. benthamiana* plants were infiltrated with a mixture of *Agrobacterium* cultures containing pBBWV2-RP1-R1 + pBBWV2-R2-53/37-Flag (this combination was designated as BBWV2-53/37-Flag) as previously described (13). At 14 days post-infiltration (dpi), crude extracts were obtained from systemic leaves of the BBWV2-53/37-Flag-infected plants by homogenizing the leaves in three volumes of protein extraction buffer (20 mM Tris–HCl at pH 7.5, 300 mM NaCl, 5 mM MgCl_2_, 5 mM dithiothreitol, 0.5% Triton X-100, proteinase inhibitor cocktail [Sigma, USA]). Cell debris was removed via centrifugation at 18,000 *g* for 20 min at 4 °C. The resulting supernatants were then incubated with anti-Flag antibody-conjugated magnetic beads (Thermo Fisher Scientific Inc., USA) for 16 h at 4 °C. After incubation, the immunocomplexes were washed five times with 1.5 mL of the protein extraction buffer. The samples were analyzed using 10% sodium dodecyl sulfate-polyacrylamide gel electrophoresis (SDS-PAGE) and stained with Coomassie blue; the Xpert prestained protein marker (GenDEPOT, USA) was used as a molecular weight marker. The bands of interest were excised from the gel for in-gel trypsin digestion followed by LC–MS/MS analysis as described previously (19). LC-MS/MS analysis was conducted at the Yonsei Proteome Research Center (Seoul, South Korea). Xcalibur (Thermo Fisher Scientific Inc., USA) was used to generate the peak lists. The data were identified by searching the National Center for Biotechnology Information database using the MASCOT search engine (http://www.matrixscience.com, Matrix Science, USA).

### Yeast two-hybrid assay (YTHA)

YTHA was performed using the MATCHMAKER Two-Hybrid System 2 (Clontech, USA) as described previously (20). The coding sequences of VP37, VP53, and HSP90 were amplified using Q5 High-Fidelity DNA polymerase (NEB, USA) and the appropriate primer pairs (Supplementary Table S1). HSP90 was fused downstream of GAL4-AD through in-frame insertion at the *Sma*I and *Xho*I sites of the pGADT7 vector. VP37 and VP53 were fused downstream of GAL4-BD via in-frame insertion using the *EcoR*I and *BamH*I sites of the pGBKT7 vector. The sequences of all constructed clones were validated using DNA sequencing. Yeast competent cells (AH109) were co-transformed with pGADT7 and pGBKT derivatives using the lithium acetate method (21). They were then selected on SD/-Leu/-Trp (SD/-LW), SD/-Leu/-Trp/-His (SD/-LWH), or SD/-Leu/-Trp/-His/-Ade (SD/-LWHA) media. The interaction between the SV40 large T antigen_(84-708)_ and murine p53_(72-390)_ was used as a positive control, whereas the interaction with human lamin C_(66-230)_ served as a negative control. All protein-protein interactions were validated by conducting α-galactosidase activity assays using the X-α-Gal reagent (Clontech, USA).

### Co-immunoprecipitation (Co-IP) and Western blot analysis

DNA templates for T7 *in vitro* transcription were prepared via PCR amplification using specific primers and templates (Supplementary Table S1): HSP90-T7-Fw and 3E-PolyA-Rv for HSP90-Flag and VP37-T7-Fw and 3E-PolyA-Rv for VP37-Myc. The resulting PCR products were purified using the GENECLEAN Turbo kit and subjected to *in vitro* coupled transcription/translation reactions using the TNT® Quick Coupled Transcription/Translation System (Promega, USA) according to the manufacturer’s instructions. The reaction products (i.e. HSP90-Flag and VP37-Myc) were mixed accordingly for Co-IP and subsequently incubated with anti-Flag antibody-conjugated magnetic beads (Thermo Fisher Scientific Inc., USA) for 1h at 26 °C. Next, the immunocomplexes were washed five times with 1.5 mL of protein extraction buffer and subjected to Western blot analysis. The resulting samples were separated through 12% SDS-PAGE and transferred onto a polyvinylidene difluoride membrane. The membrane was probed with either anti-Flag (Clontech, Japan) or anti-Myc (Santa Cruz Biotechnology, USA) primary antibodies. A secondary antibody conjugated to horseradish peroxidase (Cell Signaling Technology, USA) was used to visualize the antigens though an ECL Western blotting detection system (GE Healthcare Life Sciences, USA).

### Bimolecular fluorescence complementation assay (BiFC), ectopic expression of fluorescently tagged proteins, and cellular fluorescence imaging

The VP37 and HSP90 coding sequences were PCR-amplified from the clones used in YTHA using specific primer pairs (Supplementary Table S1) and inserted into the pENTR^TM^1A vector (Invitrogen, USA) using the *Sal*I and *Xho*I sites. The resulting clones, pENTR^TM^1A-VP37 and pENTR^TM^1A-HSP90, were then used to subclone VP37 and HSP90 into the BiFC vector pSAT4-DEST-nEYFP-C1 (containing the N-terminal half of YFP, comprising amino acids 1-174 [nYFP]) and pSAT5-DEST-cEYFP-C1 (containing the C-terminal half of YFP, comprising amino acids 175-239 [cYFP]), respectively, using the LR Clonase™ II Enzyme mix (Invitrogen, USA). The resulting constructs were named nYFP-VP37 and cYFP-HSP90, respectively. The sequences of all constructed clones were validated through DNA sequencing. PZP-YFP was used to express free YFP upon agroinfiltration (22). PZP-PDLP5-RFP was used to express PD-localizing protein 5 tagged with RFP (PDLP5-RFP), a well-known PD marker (23). PZP-ER-CFP, which expresses CFP fused with the endoplasmic reticulum (ER)-targeting signal and ER retention sequence (KDEL) (ER-CFP), was used to express an ER marker (22). PZP-VP37-GFP was used to examine the subcellular localization of VP37 (13). The coding sequence of HSP90 was PCR-amplified using the appropriate primers (Supplementary Table S1). Next, it was inserted in-frame upstream of the GFP gene in the PZP-GFP vector using the *Stu*I and *Spe*I sites (13). The resulting construct, PZP-HSP90-GFP, was used to examine the subcellular localization of HSP90. Plasmid DNA of the binary vector constructs, including the BiFC clones, was transformed into the *Agrobacterium* strain EHA105 and delivered to *N. benthamiana* leaves through agroinfiltration (22). Cellular fluorescence signals in plant leaves agroinfiltrated with BiFC and other constructs expressing fluorescent proteins were observed at 3 dpi using a Leica SP8 laser-scanning confocal microscope (Leica, Germany) equipped with specific laser/filter combinations for YFP (excitation at 514 nm), RFP (excitation at 594 nm), GFP (excitation at 488 nm), and chloroplasts (excitation at 568 nm).

### Viral sources, RNA extraction, and quantification

Full-length cDNA clones of BBWV2-RP1 and its derivative constructs were generated in previous studies (18, 24, 25). We used these viral sources and inoculated them into the leaves of 2-week-old *N. benthamiana* plants via agroinfiltration. The combination of pBBWV2-RP1-R1 and pBBWV2-R2-GFP was designated as BBWV2-GFP (26). The β-glucuronidase (GUS) gene was PCR-amplified using a specific primer pair (Supplementary Table S1). It was then cloned into pBBWV2-R2-OE (18) using the *Bgl*II and *Avr*II sites to generate pBBWV2-R2-GUS. The combination of pBBWV2-RP1-R1 and pBBWV2-R2-GUS was designated as BBWV2-GUS. RNA was extracted using the Hybrid-R^TM^ RNA extraction kit (GeneAll, South Korea) according to the manufacturer’s instructions. To confirm the systemic infection of plants inoculated with BBWV2, total RNA was isolated from the upper non-inoculated leaves and subjected to RT-PCR using a BBWV2-specific primer pair (5’-CAGAGAAGTGGTTGGTCCCGTG-3’ and 5’-ATGGGAGGCTAGTGACCTACG-3’) (13). To quantify the accumulation levels of *HSP90* mRNAs, RT followed by quantitative PCR (RT-qPCR) was performed using the diastar^TM^ onestep multiplex qRT-PCR kit (Solgent, South Korea) and Bio-Rad CFX Maestro 2.3 system (Bio-Rad, USA) with the following specific primers: qHSP90-124-Fw (5’-CGCTAGACAAGATCCGCTTTG-3’) and qHSP90-310-Rv (5’-CCTTGGTCCCTGACCTAGCA-3’) for *HSP90* detection; Nb-actin-qRT-Fw (5’-CGAGGAGCATCCAGTCCTCT-3’) and Nb-actin-qRT-Rv (5’-GTGGCTGACACCATCACCAG-3’) for actin detection. The actin gene served as an internal reference to normalize the RT-qPCR results. Three biological replicates and five technical replicates were analyzed using RT-qPCR for each sample.

### Measurement of chlorophyll contents

The relative chlorophyll content in healthy or BBWV2-infected *N. benthamiana* leaves were measured using the SPAD 502 plus chlorophyll meter (Konica Minolta, Japan), according to the manufacturer’s instructions. The SPAD readings were obtained from three different areas of each leaf, and the experiment was performed thrice with three plants in each group.

### Tobacco rattle virus (TRV)-based virus-induced gene silencing (VIGS)

A 300-bp partial fragment of *HSP90* was PCR-amplified using a specific primer pair (Supplementary Table S1). Nest, it was cloned into pTRV2 (27) using the *Sal*I site to generate pTRV2-HSP90. pTRV-GUS, which contains a partial GUS sequence was used as a negative control (28). TRV-based VIGS constructs were agroinfiltrated into the abaxial leaf surface of 2-week-old *N. benthamiana* plants.

### Imaging of fluorescence in living plants

GFP fluorescence signals in living plants were visualized using a FOBI fluorescence imaging system (NeoScience, Korea). The system was equipped with a blue light source for excitation at 470 nm and an emission filter (530 nm short-pass) that effectively removed autofluorescence signals from chlorophyll (26).

### Geldanamycin (GDA) treatment

GDA solutions were prepared at concentrations of 0, 1, and 10 μM in 1% dimethyl sulfoxide (29). GDA solutions at each concentration were syringe-infiltrated into the leaf areas previously agroinfiltrated with BiFC constructs (i.e., nYFP-VP37 + cYFP-HSP90), BBWV2-GFP, BBWV2-GUS, or PZP-VP37-GFP.

### Histochemical GUS assay

GUS expression driven by BBWV2-GUS infection was assessed using a histochemical GUS assay as previously described (30). Briefly, *N. benthamiana* leaves agroinfiltrated with BBWV2-GUS (0.005 OD_600_) were obtained at 5 dpi. Next, they were vacuum-infiltrated with the colorimetric GUS substrate 5-bromo-4-chloro-3-indoyl β-D-glucuronic acid, cyclohexylammonium salt (X-gluc) (1.2 mM) dissolved in a buffer containing 0.5 mM potassium ferricyanide, 0.5 mM potassium ferrocyanide, and 10 mM EDTA. After 12 h of incubation at 25 °C, the leaves were bleached in 70% ethanol and examined using a dissecting microscope to measure the diameters of the GUS foci.

### Reproducibility and statistical analyses

All experimental results were obtained from three independent experiments, with each experiment involving at least three plants per condition. Statistical significance between the experimental groups was evaluated using one-way ANOVA with Tukey’s HSD test (P<0.05) or a paired Student’s t-test (*P<0.05, **P<0.01). Statistical data are presented as the mean ± SD from three replicates, with each group comprising at least nine plants.

## Results

### HSP90 is a cellular interacting partner of VP37

To identify host interacting partner proteins of VP37 and VP53, we expressed VP37-Flag and VP53-Flag in *N. benthamiana* plants using the modified infectious cDNA constructs of BBWV2 (pBBWV2-RP1-R1 and pBBWV2-R2-53/37-Flag; this combination was designated as BBWV2-53/37-Flag) (Fig. 1) (13). BBWV2-53/37-Flag contains VP53 and VP37 cistrons fused in-frame with Flag, thereby expressing both VP37-Flag and VP53-Flag during viral replication (13). *N. benthamiana* plants were agroinfiltrated with BBWV2-53/37-Flag. At 14 dpi, crude plant extracts were obtained by homogenizing the upper symptomatic leaves infected with BBWV2-53/37-Flag. The crude extracts were subsequently subjected to immunoprecipitation using anti-Flag antibody-conjugated magnetic beads. The resulting products were analyzed using SDS-PAGE followed by Coomassie Blue staining (Fig. 1).

**Fig. 1.**
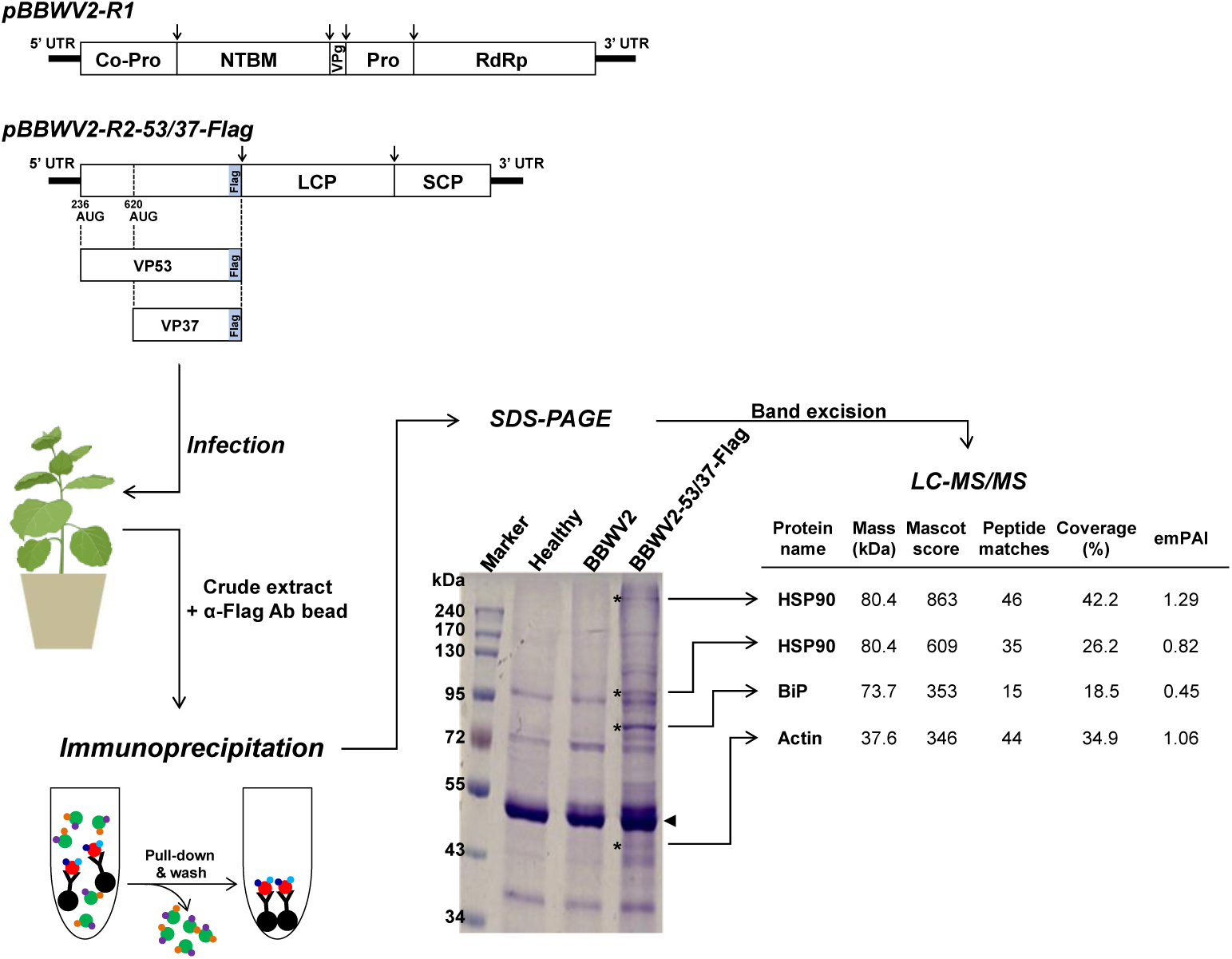
Workflow for the identification of cellular interacting protein partners of VP37 and VP53 under BBWV2 infection conditions. A schematic representation of the BBWV2 recombinant constructs shows the in-frame fusion of a Flag-tag at the C-terminus of VP37/VP53, thereby expressing VP37-Flag and VP53-Flag during viral replication. *N. benthamiana* plants were agroinfiltrated with BBWV2-53/37-Flag (a mixture of *Agrobacterium* cultures containing pBBWV2-RP1-R1 + pBBWV2-R2-53/37-Flag). At 14 dpi, crude extracts were prepared by homogenizing the upper systemically infected leaves and subjected to immunoprecipitation using magnetic beads conjugated with anti-Flag antibodies. The resulting products were analyzed using SDS-PAGE, and the bands of interest (marked with asterisks) were analyzed through LC-MS/MS. The identified proteins and their MS/MS spectral data are presented.

Four specific dominant bands in the lane containing immunoprecipitation products from leaf samples infected with BBWV2-53/37-Flag were excised from the gel and subjected to LC-MS/MS analysis. The identified proteins and their MS/MS spectral information are shown in Figure 1. The bands at positions 320 and 95 kDa were identified as HSP90. In addition, the bands at positions 80 and 45 kDa were identified as ER-luminal binding protein (BiP) and actin, respectively. The identification of HSP90 from the bands at positions 320 and 95 kDa might be attributed to its role as a molecular chaperone that interacts with diverse proteins or its capacity to form oligomers. Based on our approach, we identified three candidate host interacting partners of either VP37 or VP53. In this study, we further investigated whether HSP90 acts as a host factor by interacting with VP37 or VP53 and explored its functional role in BBWV2 infection.

We initially performed YTHA to verify whether HSP90 interacts with VP37 and/or VP53. Our results revealed that HSP90 interacts with VP37 but not VP53 under low stringency conditions (Fig. 2A). A Co-IP assay was subsequently performed to confirm the interaction between HSP90 and VP37. HSP90 and VP37, which were tagged with Flag and Myc, respectively (i.e., HSP90-Flag and VP37-Myc), were expressed using an *in vitro* translation system and subjected to Co-IP using anti-Flag antibodies. As shown in Fig. 2B, HSP90 directly interacted with VP37.

**Fig. 2.**
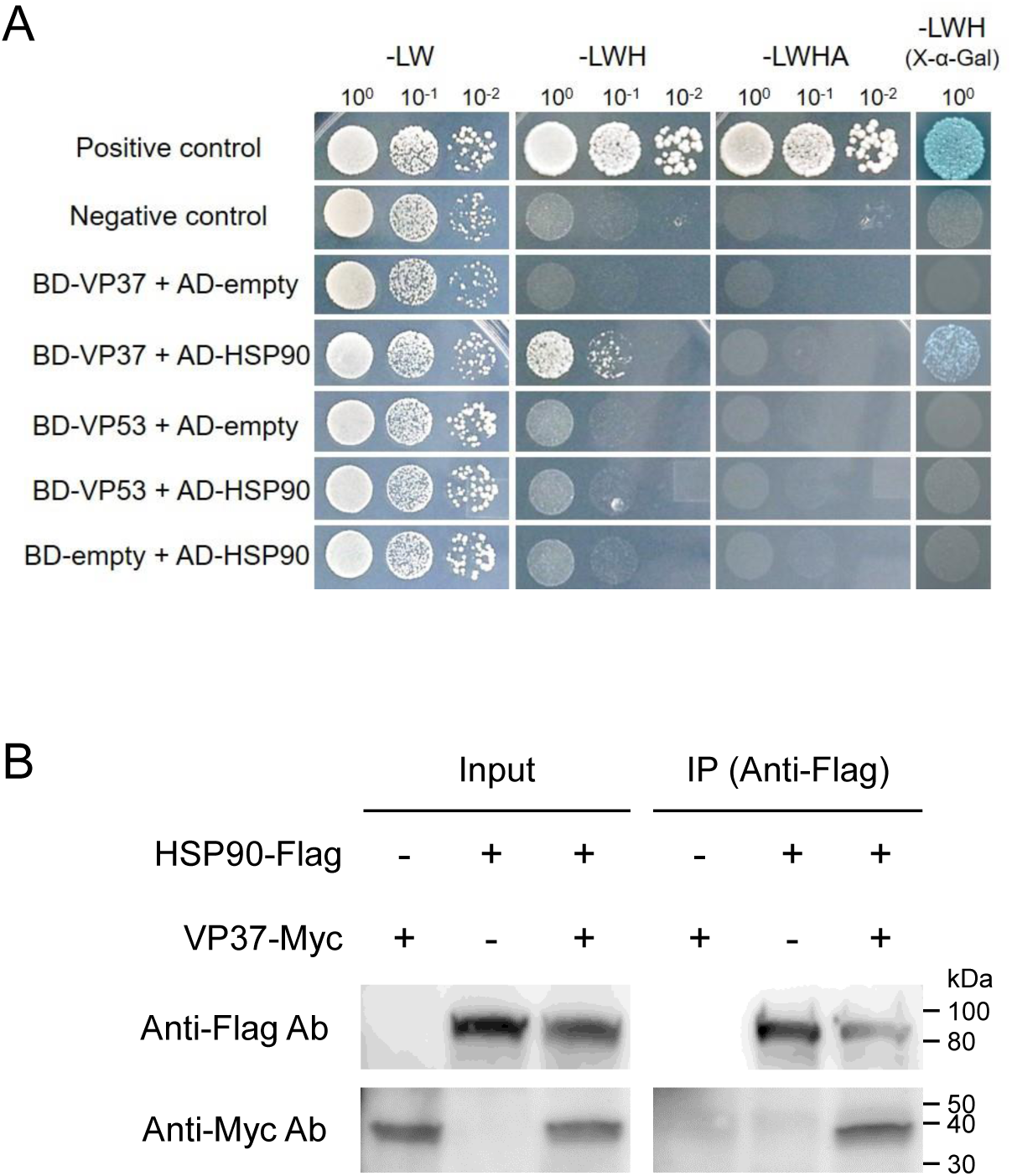
HSP90 is a cellular interacting partner of VP37. (A) The interactions between HSP90 and either VP37 or VP53 in YTHA. VP37 and VP53 were fused downstream of GAL4-BD in pGBKT7, while HSP90 was fused downstream of GAL4-AD in pGADT7. Yeast cells co-transformed with the indicated constructs were selected on SD/-LWH and SD/-LWHA agar media following gradient dilution (10^0^, 10^-1^, and 10^-2^), and their α-galactosidase activities were assessed on SD/-LWH agar media containing X-α-Gal. Interactions between the SV40 large T antigen(_84-708_) and either murine p53(_72-390_) or human lamin C_(66-230)_ were used as positive and negative controls, respectively. (B) Assessment of the HSP90-VP37 interaction through Co-IP. HSP90-Flag and VP37-Myc proteins were produced using an *in vitro* coupled transcription/translation system. The HSP90-Flag, VP37-Myc, and their combined mixture (HSP90-Flag + VP37-Myc) were immunoprecipitated using anti-Flag antibody-conjugated magnetic beads. The resulting products were analyzed by Western blotting using anti-Flag or anti-Myc antibodies.

To further confirm the HSP90-VP37 interaction *in planta* and identify the subcellular location of this interaction, we performed BiFC using an *Agrobacterium*-mediated gene expression method (22). VP37 was tagged with nYFP at the N-terminus (nYFP-VP37), whereas HSP90 was tagged with cYFP at the N-terminus (cYFP-HSP90). nYFP-VP37 and cYFP-HSP90 were co-expressed via agroinfiltration in *N. benthamiana* leaves. The reconstructed YFP signals were monitored at 3 dpi in epidermal cells using a confocal microscope. Strong YFP signals were observed as punctate spots along the cell periphery when nYFP-VP37 and cYFP-HSP90 were co-expressed (Fig. 3A). These spots were co-localized with PDLP5-RFP, which was used as a PD marker (31, 32), indicating that HSP90 and VP37 interacted at the PD (Fig. 3B).

**Fig. 3.**
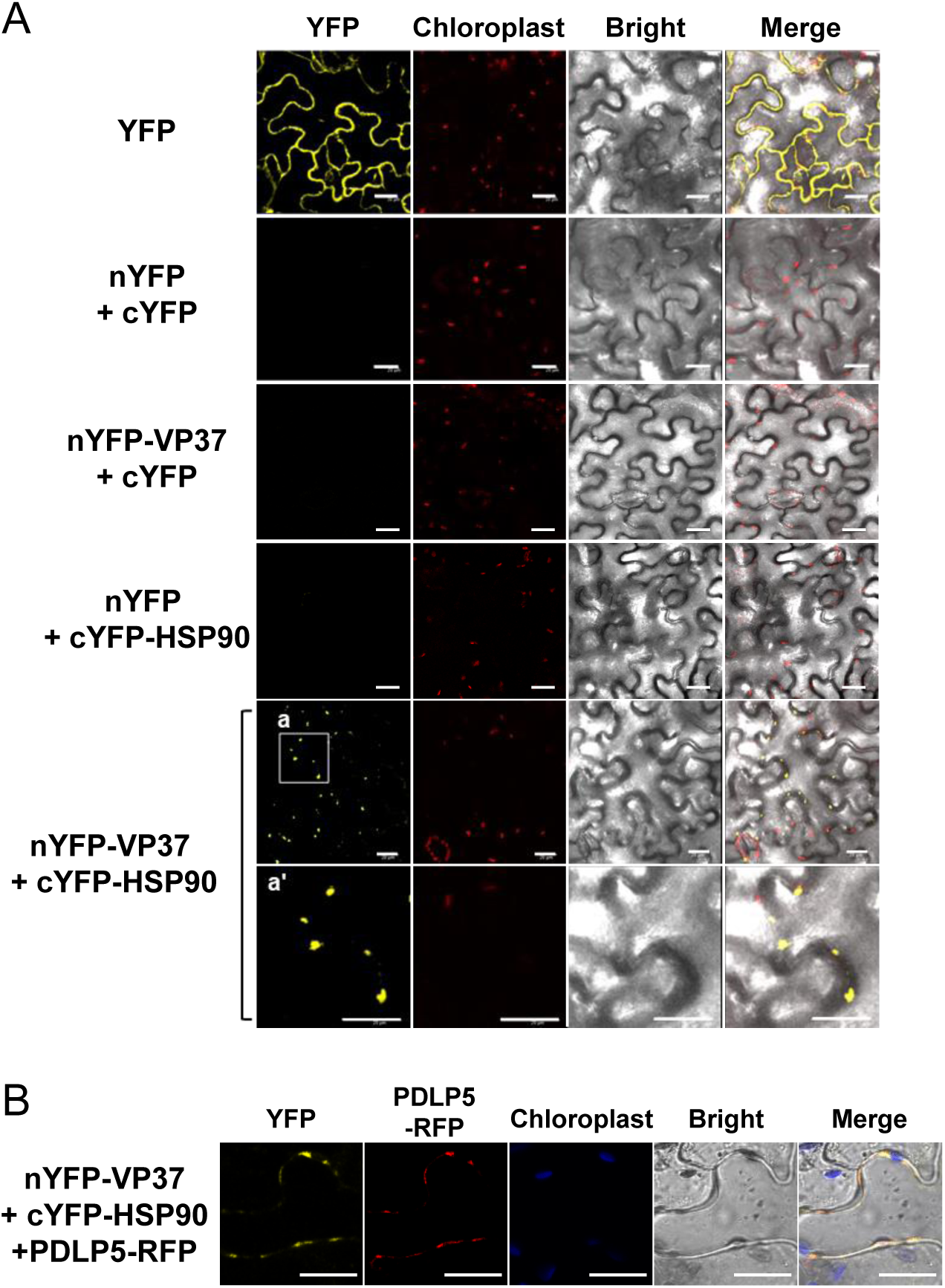
*In vivo* characterization of the HSP90-VP37 interaction using BiFC assay. (A) *In vivo* visualization of the HSP90-VP37 interaction. The recombinant proteins indicated on the left side of the panels were expressed in *N. benthamiana* leaves using an *Agrobacterium*-mediated gene expression method. The reconstructed YFP signals in the epidermal cells were observed using confocal microscopy at 3 dpi. Bar = 20μm. (B) The HSP90-VP37 interaction occurs at the PD. The indicated recombinant proteins were expressed in *N. benthamiana* leaves using an *Agrobacterium*-mediated gene expression method. PDLP5-RFP was used as a well-known PD marker. The reconstructed YFP signals in the epidermal cells were observed using confocal microscopy at 3 dpi. Bar = 20μm.

### BBWV2 infection upregulates the expression of *HSP90* and induces ER rearrangement in plant cells

HSP90 is a highly conserved molecular chaperone that plays vital roles in various cellular processes in plants, including protein homeostasis, development, signal transduction, and stress responses (33, 34). HSP90 also plays a multifaceted role in the interaction between plant viruses and their hosts (35–37). *HSP90* expression is typically upregulated in response to various stresses, including heat, drought, and pathogen attack (34). As shown in Fig. 4A, RT-qPCR analysis revealed that *HSP90* expression was upregulated upon BBWV2 infection in *N. benthamiana* plants. The upregulation of *HSP90* is strongly associated with ER stress responses (38). In addition, infection with BBWV2 (strain RP1) induced mild mosaic symptoms accompanied by a decrease in chlorophyll content in the plant leaves (Fig. 4B and C). To examine whether BBWV2 alter the ER structure in infected cells, we observed the ER structure after transiently expressing a fluorescent ER marker (ER-CFP) in cells infected with BBWV2. Confocal microscopy revealed that BBWV2 infection induced obvious ER rearrangement, forming punctate bodies along the ER network (Fig. 4D).

**Fig. 4.**
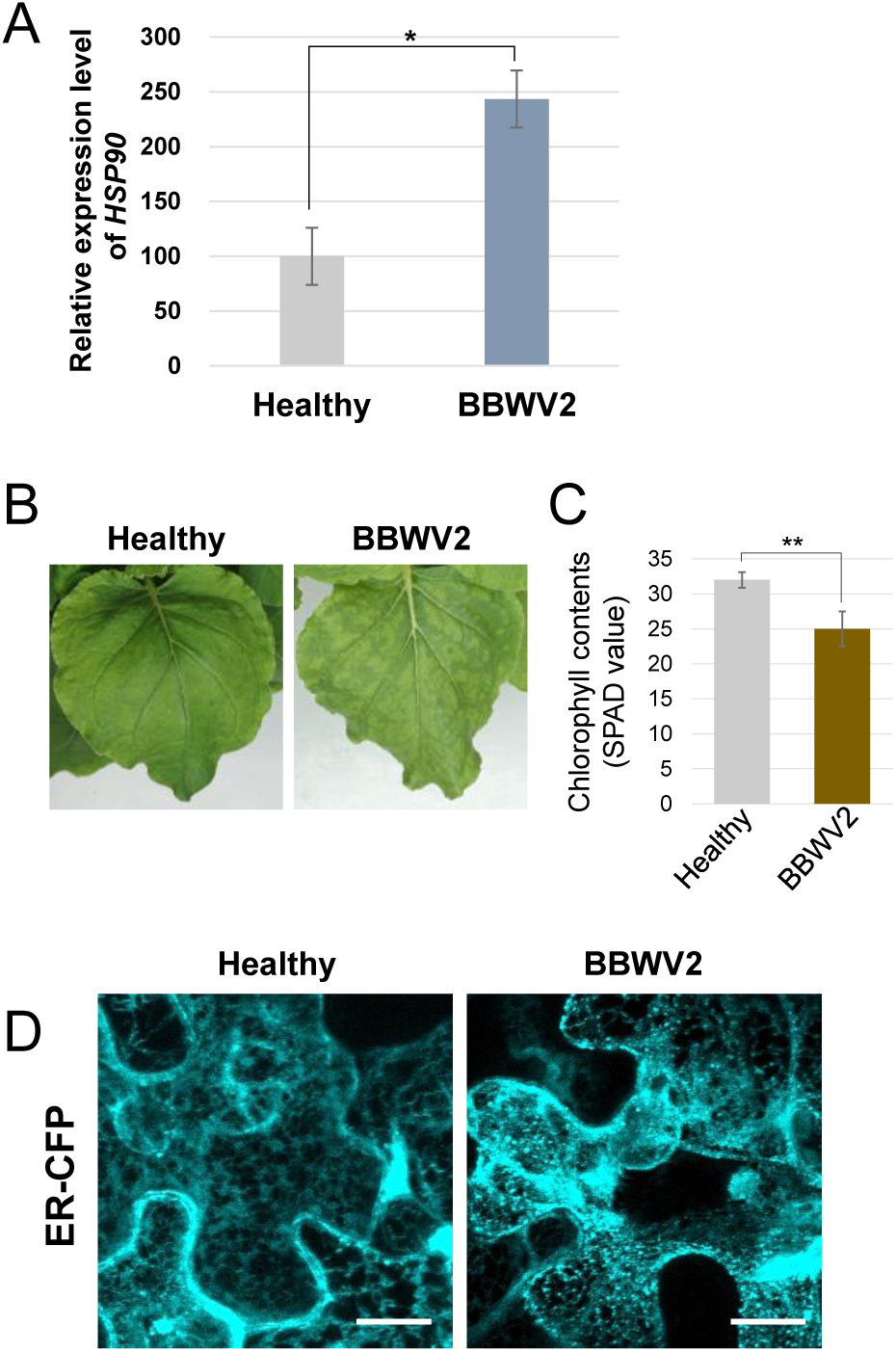
BBWV2 upregulates *HSP90* expression and induces ER rearrangement in infected plant cells. (A) BBWV2 infection upregulated *HSP90* expression. Total RNA was isolated from the upper symptomatic leaves of *N. benthamiana* plants infected with BBWV2 at 12 dpi, and the relative accumulation levels of *HSP90* were analyzed by RT-qPCR. Data are presented as the mean ± SD of three replicates, with each group containing nine plants. Statistical significance was determined using a paired Student’s t-test: \**P*<0.05. (B) *N. benthamiana* leaves infected with BBWV2 exhibited mosaic symptoms. *N. benthamiana* plants were agroinfiltrated with pBBWV2-RP1, and symptoms on the upper uninoculated leaves were observed at 12 dpi. (C) BBWV2 infection caused a decrease in chlorophyll contents. The relative chlorophyll contents in healthy or BBWV2-infected leaves of *N. benthamiana* plants were assessed by measuring SPAD values. Data are presented as the mean ± SD of three replicates, with each group containing nine plants. Statistical significance was determined using a paired Student’s t-test: \*\**P*<0.01. (D) BBWV2 infection induced ER rearrangement. ER-CFP, an ER marker, was expressed in healthy and BBWV2-infected leaves of *N. benthamiana* plants using an *Agrobacterium*-mediated gene expression method. Subcellular fluorescence signals in the epidermal cells were observed using confocal microscopy at 3 dpi. Bar = 10μm.

### HSP90 is crucial for systemic infection of BBWV2

To examine the requirement of HSP90 for BBWV2 infection, we first performed a loss-of-function analysis of *HSP90* using a TRV-based VIGS system (39). *N. benthamiana* plants inoculated with a TRV construct containing a partial sequence of GUS (pTRV-GUS) were used as negative controls (28). Consistent with previous studies (40, 41), *HSP90* silencing resulted in strong growth inhibition and leaf yellowing in *N. benthamiana* plants (Fig. 5A and Supplementary Fig. S1). To evaluate the silencing efficiency of *HSP90*, RT-qPCR was performed using RNA extracted from the systemic leaves of inoculated *N. benthamiana* plants. The result confirmed the efficient silencing of *HSP90* (Fig. 5B and Supplementary Fig. S2). One week after inoculation with the TRV-based VIGS constructs, the plants were agroinfiltrated with BBWV2-GFP (a mixture of *Agrobacterium* cultures containing pBBWV2-RP1-R1 + pBBWV2-R2-GFP), enabling visual tracking of BBWV2 infection dynamics in living plants (18, 26). The inoculated plants were observed using a FOBI fluorescence imaging system starting at 4 dpi with BBWV2-GFP. Although BBWV2-GFP successfully established systemic infection in the plants inoculated with mock or pTRV-GUS, BBWV2 infection was significantly inhibited in *HSP90*-silenced plants (Fig. 5C). Our results suggest that HSP90 is a host cellular protein essential for BBWV2 systemic infection.

**Fig. 5.**
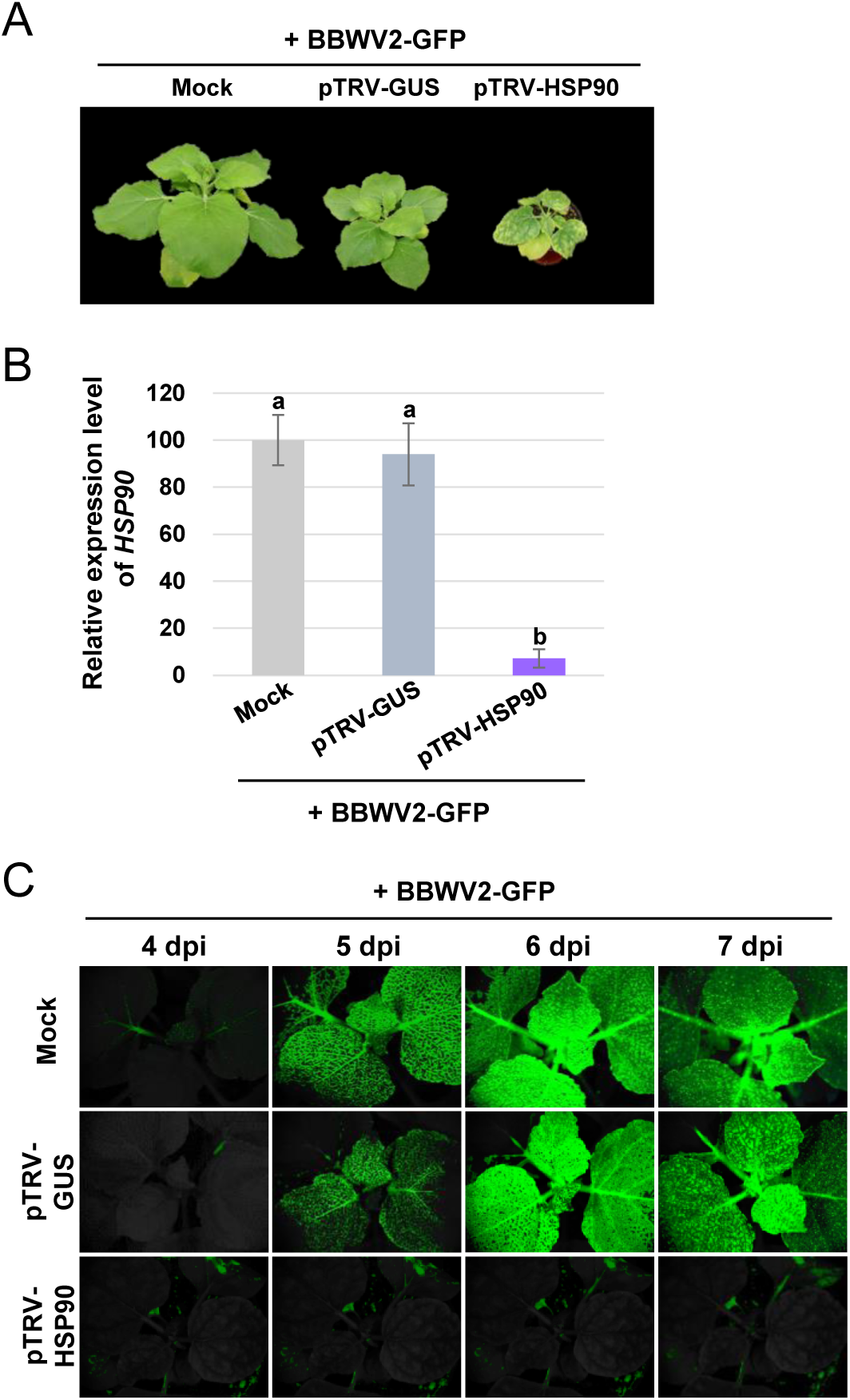
Effects of *HSP90* silencing on the systemic spread of BBWV2. 2-week-old *N. benthamiana* plants agroinfiltrated with mock (*Agrobacterium* containing no binary vector), pTRV-GUS, or pTRV-GUS. One week later, the plants were agroinfiltrated with BBWV2-GFP. The plants inoculated with mock or pTRV-GUS served as negative controls. (A) Phenotypes of BBWV2-infected *N. benthamiana* plants following *HSP90* silencing. *HSP90* silencing caused strong growth inhibition and leaf yellowing in BBWV2-infeced *N. benthamiana* plants. The plants were photographed at 14 dpi with BBWV2-GFP. (B) RT-qPCR analysis of *HSP90* expression. Total RNA isolated from the systemic leaves of inoculated *N. benthamiana* plants at 14 dpi was analyzed using RT-qPCR to assess the silencing efficiency of *HSP90*. (C) Time-course observation of the systemic spread of BBWV2-GFP in *HSP90*-silenced *N. benthamiana* plants. The plants agroinfiltrated with BBWV2-GFP were observed using a FOBI fluorescence imaging system at 4, 5, 6, and 7 dpi. Data shown are representatives of at least three independent experiments.

### Chaperone function of HSP90 is required for its interaction with VP37 and cell-to-cell movement of BBWV2

The chaperone function of HSP90 is tightly regulated by the ATPase activity presented by its N-terminal domain (42, 43). The function of HSP90 can be inhibited by GDA, which blocks the ATPase activity of HSP90 (44). To examine whether the chaperone function of HSP90 is required for its interaction with VP37, we performed additional BiFC experiments under GDA treatment conditions. nYFP-VP37 and cYFP-HSP90 were co-expressed via agroinfiltration in *N. benthamiana* leaves. After 36 h, the agroinfiltrated leaf areas were syringe-infiltrated with 0, 1, or 10 µM GDA. The interaction between nYFP-VP37 and cYFP-HSP90 was then monitored at 3 dpi using confocal microscopy. As shown in Fig. 6, strong YFP signals were observed as punctate spots in the absence of GDA. However, no YFP signals were detected following treatment with 1 or 10 µM GDA (Fig. 6). In addition, GDA treatment did not induce significant changes in the expression or subcellular localization patterns of HSP90 (Supplementary Fig. S2). These results suggest that the ATP-dependent chaperone function of HSP90 is essential for the HSP90-VP37 interaction.

**Fig. 6.**
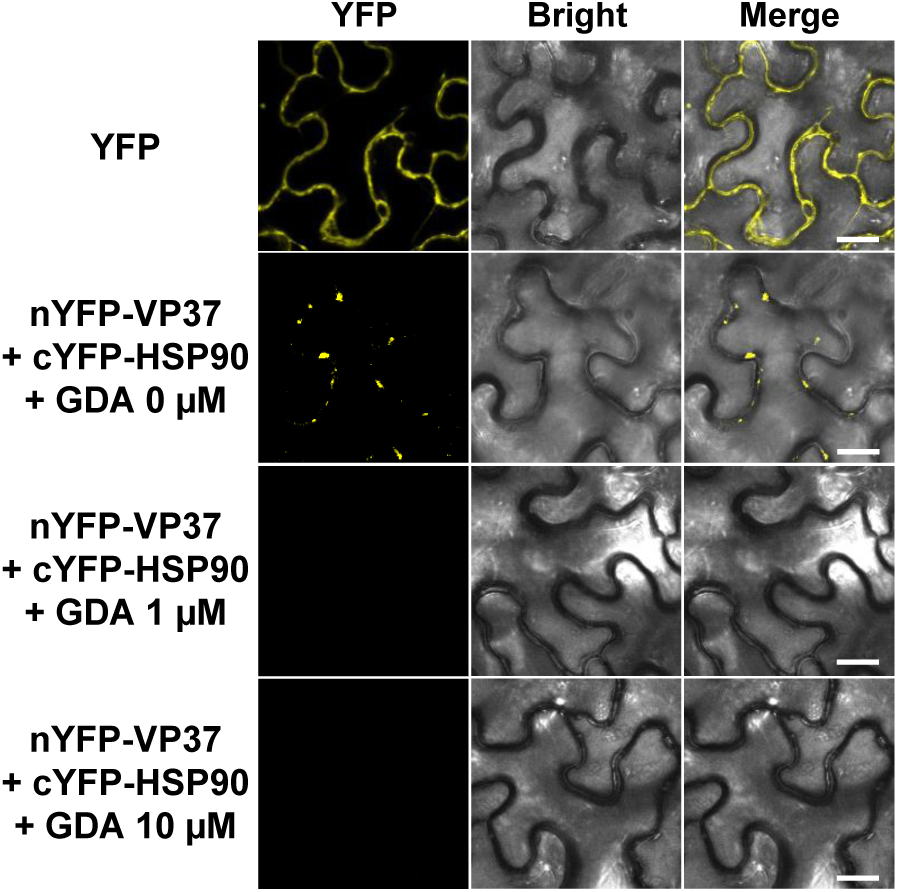
Effects of inhibiting the chaperone function of HSP90 on the HSP90-VP37 interaction. nYFP-VP37 and cYFP-HSP90 were co-expressed in *N. benthamiana* leaves using an *Agrobacterium*-mediated gene expression method. After 36 h, 0, 1, or 10 µM GDA was syringe-infiltrated into the agroinfiltrated leaf area. The reconstructed YFP signals in the epidermal cells were observed using confocal microscopy at 3 dpi. Bar = 20μm.

We further examined whether the chaperone function of HSP90 is required for the systemic spread of BBWV2. *N. benthamiana* leaves were agroinfiltrated with BBWV2-GFP. After 36 h, the agroinfiltrated leaf areas were syringe-infiltrated with 0 or 10 µM GDA. The plants were observed using a FOBI fluorescence imaging system starting at 5.5 dpi with BBWV2-GFP. Although strong GFP signals were observed as early as 5.5 dpi in plants treated with 0 µM GDA, clear GFP signals began to appear from 6.5 dpi in those treated with 10 µM GDA (Fig. 7A). This indicated that BBWV2 movement was suppressed in the GDA-treated leaf area, consequently delaying the systemic infection of BBWV2.

**Fig. 7.**
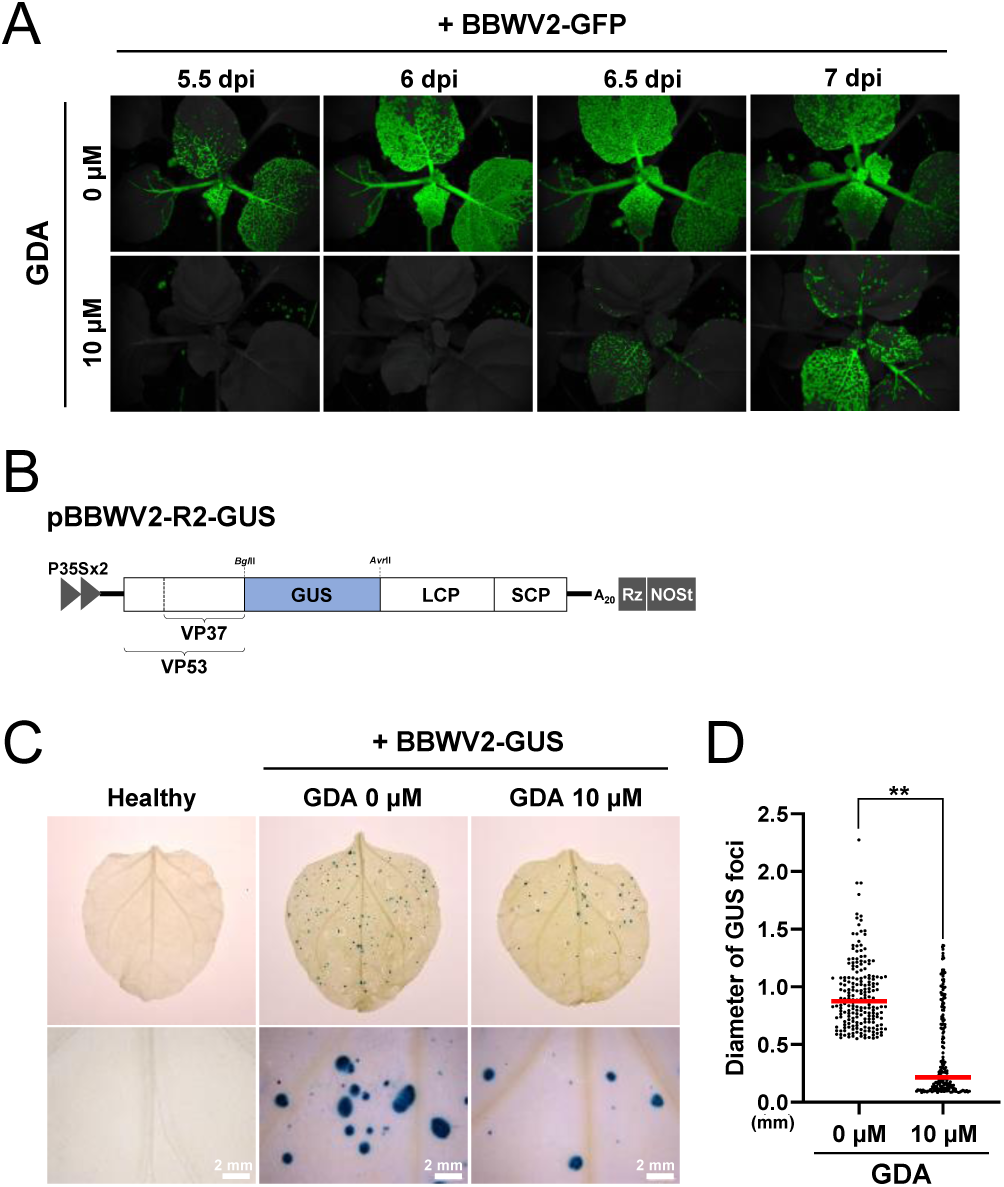
Chaperone function of HSP90 is required for the systemic spread and cell-to-cell movement of BBWV2. (A) Effects of inhibiting the chaperone function of HSP90 on the systemic spread of BBWV2. *N. benthamiana* plants agroinfiltrated with BBWV2-GFP. After 36 h, 0 or 10 µM GDA was syringe-infiltrated into the agroinfiltrated leaf area. The plants were observed using a FOBI fluorescence imaging system at 5.5, 6, 6.5, and 7 dpi. Data shown are representatives of at least three independent experiments. (B) Schematic representation of the BBWV2 RNA2 construct tagged with GUS (pBBWV2-R2-stagRFP), designed to express GUS during viral replication. (C) Effects of inhibiting the chaperone function of HSP90 on the cell-to-cell movement of BBWV2. *N. benthamiana* plants were agroinfiltrated with BBWV2-GUS. After 36 h, 0 or 10 µM GDA was syringe-infiltrated into the agroinfiltrated leaf area. The infiltrated leaves were subjected to a histochemical GUS assay at 5 dpi with BBWV2-GUS. The GUS foci were photographed at 5 dpi. Data shown are representatives of at least three independent experiments. (D) Statistical analysis of GUS foci diameters shown in panel C. The diameters of the 200 largest GUS foci were measured for each treatment across three independent experiments. The dot plot illustrates the distribution of the GUS foci diameters in ascending order, with the red horizontal bar in each group representing the mean diameter of the measured GUS foci. Statistical significance was determined using a paired Student’s t-test: \*\**P*<0.01.

We further examined whether the inhibition of HSP90 activity by GDA treatment affects the cell-to-cell movement of BBWV2. *N. benthamiana* leaves were agroinfiltrated with BBWV2-GUS (a mixture of *Agrobacterium* cultures containing pBBWV2-RP1-R1 + pBBWV2-R2-GUS) (Fig. 7B), enabling visual tracking of the cell-to-cell movement of BBWV2. After 36 h, the agroinfiltrated leaf areas were syringe-infiltrated with 0 or 10 µM GDA. At 5 dpi with BBWV2-GUS, the infiltrated leaves were subjected to a histochemical GUS assay to visually measure the extent of viral cell-to-cell movement. BBWV2-GUS spread radially to form infection foci with an average diameter of approximately 930 ± 297.3 µm following treatment with 0 µM GDA (Fig. 7C and D). In contrast, the cell-to-cell movement of BBWV2-GUS was significantly suppressed when treated with 10 µM GDA; the average diameter of foci was approximately 415.2 ± 375.3 µm (Fig. 7C and D). These results suggest that the chaperone function of HSP90 is required for the cell-to-cell movement of BBWV2.

### HSP90-VP37 interaction is required for VP37-derived tubule formation at the PD

GFP tagging at the C-terminus of VP37 (VP37-GFP) does not impair its function as a viral movement protein (13, 14). In addition, VP37-GFP is localized to the PD and forms tubule structures when expressed transiently in plant cells (13, 14). As we revealed that the HSP90-VP37 interaction can be inhibited by GDA treatment (Fig. 6), we used GDA treatment to investigate whether this interaction is essential for VP37-derived tubule formation in the PD. VP37-GFP was transiently expressed via agroinfiltration in *N. benthamiana* leaves. After 36 h, the agroinfiltrated leaf areas were syringe-infiltrated with 0 or 10 µM GDA. Next, the subcellular localization of VP37-GFP was observed at 3 dpi using confocal microscopy. In the absence of GDA treatment (0 µM), VP37-GFP predominantly accumulated as punctate spots that were positioned opposite each other on the plasma membranes of two adjacent cells, suggesting the formation of VP37-derived tubules through the PD (Fig. 8). However, when the HSP90-VP37 interaction was inhibited by treatment with 10 µM GDA, VP53-GFP punctate spots were observed only on the plasma membrane of a single cell, with no corresponding spots on the adjacent cell, suggesting that tubule formation through the PD might be impaired (Fig. 8). Our results suggest that VP37-derived tubule formation through the PD likely requires interaction with HSP90.

**Fig. 8.**
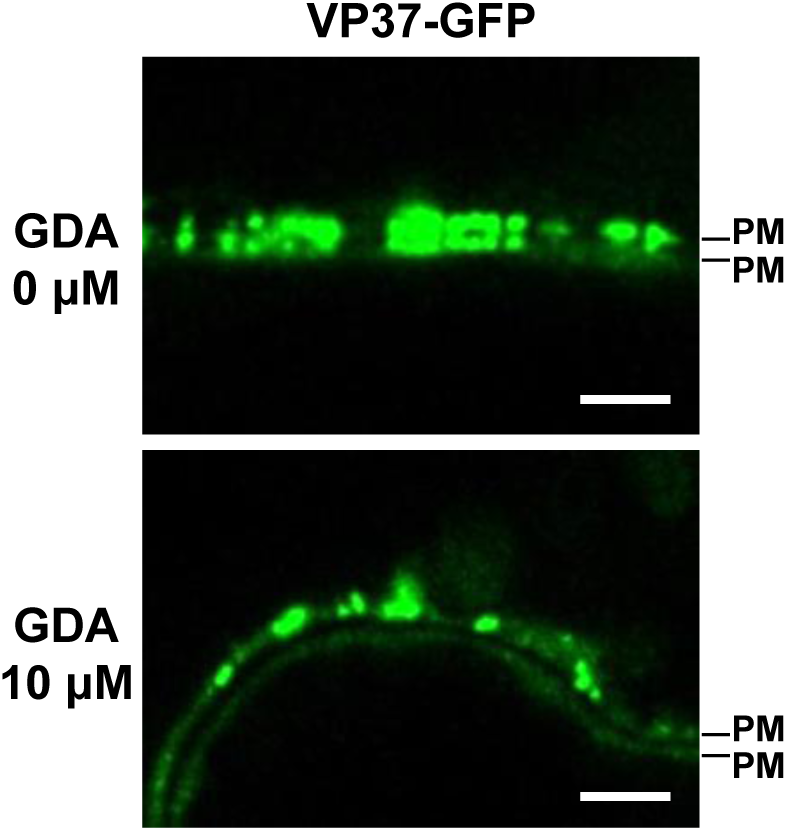
Effects of inhibiting the chaperoning function of HSP90 on VP37-derived tubule formation at PD. VP37-GFP was expressed in *N. benthamiana* leaves using an *Agrobacterium*-mediated gene expression method. After 36 h, 0 or 10 µM GDA was syringe-infiltrated into the agroinfiltrated leaf area. The subcellular localization of VP37-GFP was observed using confocal microscopy at 3 dpi. PM, plasma membrane. Bar = 10 μm.

## Discussion

During infection, viruses alter the cellular environment of their hosts and utilizes various host factors, including cellular proteins (1, 3). In addition, successful systemic infection by plant viruses relies on their ability to move from cell to cell via the PD (8, 9). Interactions between viral and host cellular proteins also play a pivotal role in viral cell-to-cell movement (6). In this study, we identified HSP90 as a host factor that interacts with VP37, the MP of BBWV2. We also demonstrated the importance of this interaction in the cell-to-cell movement and systemic infection of BBWV2.

Although few host proteins have been identified to interact with viral MPs at the PD, pectin methylesterase (PME) is a well-characterized example of a host plasmodesmal protein known to interact with the MPs of tobacco mosaic virus, cauliflower mosaic virus, and turnip vein clearing virus (10, 45, 46). Because PME is primarily present in the cell wall around the PD, it may serve as a general peripheral target for viral MPs in PD localization (45, 47). In addition, the family of PD-located proteins (PDLPs) interacts with MPs of grapevine fanleaf virus (GFLV) and cauliflower mosaic virus (CaMV) at the PD, facilitating the movement of these viruses (31). In PDLP knock-out plants, tubule formation at the PD and systemic infection with GFLV and CaMV were significantly suppressed (31).

HSP90 is a multifunctional chaperone crucial for protein homeostasis, particularly assisting in the refolding or degradation of misfolded or aggregated proteins that arise from various stress responses (34). In the context of viral infection, HSP90 is involved in the assembly of the replication complexes of various RNA viruses, including bamboo mosaic virus (BaMV) and red clover necrotic mosaic virus (RCNMV) (35, 36). In BaMV, HSP90 facilitates the proper folding of BaMV replicase and recruitment of the viral RNA template during the early stages of replication (36). In RCNMV, HSP90 is essential for viral replication and interacts with p27, a virus-encoded replicase component (35). The infectivity of BaMV and RCNMV was significantly reduced following *HSP90* silencing or GDA treatment in *N. benthamiana* plants (35, 36). However, the role of HSP90 in viral transport remains largely understudied.

In this study, VIGS and GDA treatment experiments demonstrated that the HSP90 is required for the systemic spread of BBWV2 (Figs. 5 and 7), indicating that it is a crucial host factor for BBWV2 infectivity. In addition, HSP90 interacted with VP37 at the PD (Fig. 3). Interestingly, the formation of VP37-derived tubules through the PD appeared to be impaired when the HSP90-VP37 interaction was inhibited by GDA treatment (Fig. 8). We hypothesized that the inhibition of BBWV2 cell-to-cell movement after GDA treatment may result from impaired tubule formation. These findings emphasize the requirement of the chaperone function of HSP90 in the HSP90-VP37 interaction and BBWV2 cell-to-cell movement, as GDA treatment disrupts the ATP-dependent chaperone activity of HSP90 (44). VP37 forms tubule structures specifically at the PD, but not in the cytoplasm (13, 14, 16). Moreover, the HSP90-VP37 interaction exclusively occurs at the PD (Fig. 3). These observations suggest that the chaperone activity of HSP90 may induce a conformational change in VP37, thereby facilitating the formation of VP37-derived tubules through the PD. Interestingly, a previous study revealed that a significant number of plant viral MPs exhibits unexpected amino acid sequence homology with HSP90 (48). These structural similarities between various plant viral MPs and HSP90 suggest a potential requirement for the functional activities of HSP90, such as its molecular chaperone activity, in facilitating viral cell-to-cell movement. In addition to promoting stable protein folding, molecular chaperones play key roles in the assembly of multimeric protein complexes (49–52). Therefore, it is possible that chaperone activity is involved in the formation of virus-induced intracellular structures, such as tubules, or in the modification of the PD structure to increase its size exclusion limit.

In conclusion, HSP90 specifically interacts with VP37 at the PD, and this interaction is likely essential for the formation of VP37-derived tubules that facilitate viral transport through the PD. Our study further demonstrated that the ATP-dependent chaperone activity of HSP90, which can be disrupted by GDA treatment, is integral to this process, highlighting the important role of molecular chaperones in viral movement. Our findings provide new insights into the molecular mechanisms underlying the cell-to-cell movement of plant viruses.

## Data Availability Statement

The original contributions presented in the study are included in the article/Supplementary Material, further inquiries can be directed to the corresponding author.

## Author Contributions

JKS designed the experiments and supervised the project; MHK, SYJ, JSC, SK, YL, SP, and SJK performed the experiments; MHK, SYJ, SJK, and JKS analyzed the data; MHK and JKS wrote and revised the manuscript.

## References

1. Pallas V, Garcia JA. 2011. How do plant viruses induce disease? Interactions and interference with host components. J Gen Virol 92:2691–2705.

2. McLeish MJ, Fraile A, Garcia-Arenal F. 2019. Evolution of plant-virus interactions: host range and virus emergence. Curr Opin Virol 34:50–55.

3. Culver JN, Padmanabhan MS. 2007. Virus-induced disease: altering host physiology one interaction at a time. Annu Rev Phytopathol 45:221–243.

4. Wang A. 2015. Dissecting the molecular network of virus-plant interactions: the complex roles of host factors. Annu Rev Phytopathol 53:45–66.

5. Nagy PD. 2008. Yeast as a model host to explore plant virus-host interactions. Annu Rev Phytopathol 46:217–242.

6. Harries P, Ding B. 2011. Cellular factors in plant virus movement: at the leading edge of macromolecular trafficking in plants. Virology 411:237–243.

7. Lucas WJ. 2006. Plant viral movement proteins: agents for cell-to-cell trafficking of viral genomes. Virology 344:169–184.

8. Wang A. 2021. Cell-to-cell movement of plant viruses via plasmodesmata: a current perspective on potyviruses. Curr Opin Virol 48:10–16.

9. Ueki S, Citovsky V. 2011. To gate, or not to gate: regulatory mechanisms for intercellular protein transport and virus movement in plants. Mol Plant 4:782–793.

10. Boevink P, Oparka KJ. 2005. Virus-host interactions during movement processes. Plant Physiol 138:1815–1821.

11. Brown SL, Garrison DJ, May JP. 2021. Phase separation of a plant virus movement protein and cellular factors support virus-host interactions. PLoS Pathog 17:e1009622.

12. Kim MH, Kwak HR, Choi B, Kwon SJ, Seo JK. 2021. Genetic plasticity in RNA2 is associated with pathogenic diversification of broad bean wilt virus 2. Virus Res 304:198533.

13. Kim MH, Choi B, Jang SY, Choi JS, Kim S, Lee Y, Park S, Kwon SJ, Kang JH, Seo JK. 2024. The VP53 protein encoded by RNA2 of a fabavirus, broad bean wilt virus 2, is essential for viral systemic infection. Commun Biol 7:462.

14. Liu C, Ye L, Lang G, Zhang C, Hong J, Zhou X. 2011. The VP37 protein of Broad bean wilt virus 2 induces tubule-like structures in both plant and insect cells. Virus Res 155:42–47.

15. Liu C, Meng C, Xie L, Hong J, Zhou X. 2009. Cell-to-cell trafficking, subcellular distribution, and binding to coat protein of Broad bean wilt virus 2 VP37 protein. Virus Res 143:86–93.

16. Xie L, Shang W, Liu C, Zhang Q, Sunter G, Hong J, Zhou X. 2016. Mutual association of Broad bean wilt virus 2 VP37-derived tubules and plasmodesmata obtained from cytological observation. Sci Rep 6:21552.

17. Qi YJ, Zhou XP, Huang XZ, Li GX. 2002. In vivo accumulation of Broad bean wilt virus 2 VP37 protein and its ability to bind single-stranded nucleic acid. Arch Virol 147:917–928.

18. Choi B, Kwon SJ, Kim MH, Choe S, Kwak HR, Kim MK, Jung C, Seo JK. 2019. A Plant Virus-Based Vector System for Gene Function Studies in Pepper. Plant Physiol 181:867–880.

19. Seo JK, Choi HS, Kim KH. 2016. Engineering of soybean mosaic virus as a versatile tool for studying protein-protein interactions in soybean. Sci Rep 6:22436.

20. Seo JK, Kang SH, Seo BY, Jung JK, Kim KH. 2010. Mutational analysis of interaction between coat protein and helper component-proteinase of involved in aphid transmission. Mol Plant Pathol 11:265–276.

21. Schiestl RH, Gietz RD. 1989. High-Efficiency Transformation of Intact Yeast-Cells Using Single Stranded Nucleic-Acids as a Carrier. Curr Genet 16:339–346.

22. Seo JK, Kwon SJ, Rao AL. 2012. A physical interaction between viral replicase and capsid protein is required for genome-packaging specificity in an RNA virus. J Virol 86:6210–6221.

23. Kumar R, Iswanto ABB, Kumar D, Shuwei W, Oh K, Moon J, Son GH, Oh ES, Vu MH, Lee J, Lee KW, Oh MH, Kwon C, Chung WS, Kim JY, Kim SH. 2024. C-Type LECTIN receptor-like kinase 1 and ACTIN DEPOLYMERIZING FACTOR 3 are key components of plasmodesmata callose modulation. Plant Cell Environ 1–17.

24. Seo JK, Kwon SJ, Choi HS, Kim KH. 2009. Evidence for alternate states of Cucumber mosaic virus replicase assembly in positive- and negative-strand RNA synthesis. Virology 383:248–260.

25. Seo JK, Kwak HR, Choi B, Han SJ, Kim MK, Choi HS. 2017. Movement protein of broad bean wilt virus 2 serves as a determinant of symptom severity in pepper. Virus Res 242:141–145.

26. Kwon MJ, Kwon SJ, Kim MH, Choi B, Byun HS, Kwak HR, Seo JK. 2023. Visual tracking of viral infection dynamics reveals the synergistic interactions between cucumber mosaic virus and broad bean wilt virus 2. Sci Rep 13:7261.

27. Bachan S, Dinesh-Kumar SP. 2012. Tobacco rattle virus (TRV)-based virus-induced gene silencing. Methods Mol Biol 894:83–92.

28. Choe S, Choi B, Kang JH, Seo JK. 2021. Tolerance to tomato yellow leaf curl virus in transgenic tomato overexpressing a cellulose synthase-like gene. Plant Biotechnol J 19:657–659.

29. Chong LP, Wang Y, Gad N, Anderson N, Shah B, Zhao R. 2015. A highly charged region in the middle domain of plant endoplasmic reticulum (ER)-localized heat-shock protein 90 is required for resistance to tunicamycin or high calcium-induced ER stresses. J Exp Bot 66:113–124.

30. Dolja VV, McBride HJ, Carrington JC. 1992. Tagging of plant potyvirus replication and movement by insertion of beta-glucuronidase into the viral polyprotein. Proc Natl Acad Sci 89:10208–10212.

31. Amari K, Boutant E, Hofmann C, Schmitt-Keichinger C, Fernandez-Calvino L, Didier P, Lerich A, Mutterer J, Thomas CL, Heinlein M, Mely Y, Maule AJ, Ritzenthaler C. 2010. A family of plasmodesmal proteins with receptor-like properties for plant viral movement proteins. PLoS Pathog 6:e1001119.

32. Iswanto ABB, Vu MH, Shon JC, Kumar R, Wu S, Kang H, Kim DR, Son GH, Kim WY, Kwak YS, Liu KH, Kim SH, Kim JY. 2024. alpha1-COP modulates plasmodesmata function through sphingolipid enzyme regulation. J Integr Plant Biol 00:1–19.

33. Sangster TA, Queitsch C. 2005. The HSP90 chaperone complex, an emerging force in plant development and phenotypic plasticity. Curr Opin Plant Biol 8:86–92.

34. Xu ZS, Li ZY, Chen Y, Chen M, Li LC, Ma YZ. 2012. Heat shock protein 90 in plants: molecular mechanisms and roles in stress responses. Int J Mol Sci 13:15706–15723.

35. Mine A, Hyodo K, Tajima Y, Kusumanegara K, Taniguchi T, Kaido M, Mise K, Taniguchi H, Okuno T. 2012. Differential roles of Hsp70 and Hsp90 in the assembly of the replicase complex of a positive-strand RNA plant virus. J Virol 86:12091–12104.

36. Huang YW, Hu CC, Liou MR, Chang BY, Tsai CH, Meng M, Lin NS, Hsu YH. 2012. Hsp90 interacts specifically with viral RNA and differentially regulates replication initiation of Bamboo mosaic virus and associated satellite RNA. PLoS Pathog 8:e1002726.

37. Gorovits R, Czosnek H. 2017. The Involvement of Heat Shock Proteins in the Establishment of Tomato Yellow Leaf Curl Virus Infection. Front Plant Sci 8:355.

38. Manghwar H, Li J. 2022. Endoplasmic Reticulum Stress and Unfolded Protein Response Signaling in Plants. Int J Mol Sci 23:828.

39. Liu Y, Schiff M, Dinesh-Kumar SP. 2002. Virus-induced gene silencing in tomato. Plant J 31:777–786.

40. Shibata Y, Kawakita K, Takemoto D. 2011. SGT1 and HSP90 are essential for age-related non-host resistance of Nicotiana benthamiana against the oomycete pathogen Phytophthora infestans. Physiol Mol Plant Pathol 75:120–128.

41. Xu C, Guo H, Li R, Lan X, Zhang Y, Xie Q, Zhu D, Mu Q, Wang Z, An M, Xia Z, Wu Y. 2023. Transcriptomic and functional analyses reveal the molecular mechanisms underlying Fe-mediated tobacco resistance to potato virus Y infection. Front Plant Sci 14:1163679.

42. Obermann WM, Sondermann H, Russo AA, Pavletich NP, Hartl FU. 1998. In vivo function of Hsp90 is dependent on ATP binding and ATP hydrolysis. J Cell Biol 143:901–910.

43. Siligardi G, Hu B, Panaretou B, Piper PW, Pearl LH, Prodromou C. 2004. Co-chaperone regulation of conformational switching in the Hsp90 ATPase cycle. J Biol Chem 279:51989–51998.

44. Stebbins CE, Russo AA, Schneider C, Rosen N, Hartl FU, Pavletich NP. 1997. Crystal structure of an Hsp90-geldanamycin complex: targeting of a protein chaperone by an antitumor agent. Cell 89:239–250.

45. Chen MH, Sheng JS, Hind G, Handa AK, Citovsky V. 2000. Interaction between the tobacco mosaic virus movement protein and host cell pectin methylesterases is required for viral cell-to-cell movement. EMBO J 19:913–920.

46. Dorokhov YL, Makinen K, Frolova OY, Merits A, Saarinen J, Kalkkinen N, Atabekov JG, Saarma M. 1999. A novel function for a ubiquitous plant enzyme pectin methylesterase: the host-cell receptor for the tobacco mosaic virus movement protein. FEBS Lett 461:223–228.

47. Morvan O, Quentin M, Jauneau A, Mareck A, Morvan C. 1998. Immunogold localization of pectin methylesterases in the cortical tissues of flax hypocotyl. Protoplasma 202:175–184.

48. Koonin EV, Mushegian AR, Ryabov EV, Dolja VV. 1991. Diverse groups of plant RNA and DNA viruses share related movement proteins that may possess chaperone-like activity. J Gen Virol 72:2895–2903.

49. Saibil H. 2013. Chaperone machines for protein folding, unfolding and disaggregation. Nat Rev Mol Cell Biol 14:630–642.

50. Seo HW, Seo JP, Jung G. 2018. Heat shock protein 70 and heat shock protein 90 synergistically increase hepatitis B viral capsid assembly. Biochem Biophys Res Commun 503:2892–2898.

51. Ellis RJ. 2006. Molecular chaperones: assisting assembly in addition to folding. Trends Biochem Sci 31:395–401.

52. Li K, Jiang Q, Bai X, Yang YF, Ruan MY, Cai SQ. 2017. Tetrameric Assembly of K(+) Channels Requires ER-Located Chaperone Proteins. Mol Cell 65:52–65.

