## Supplementary data for "HSP90 interacts with VP37 to facilitate the cell-to-cell movement of broad bean wilt virus 2"

### Supplementary Figure S1

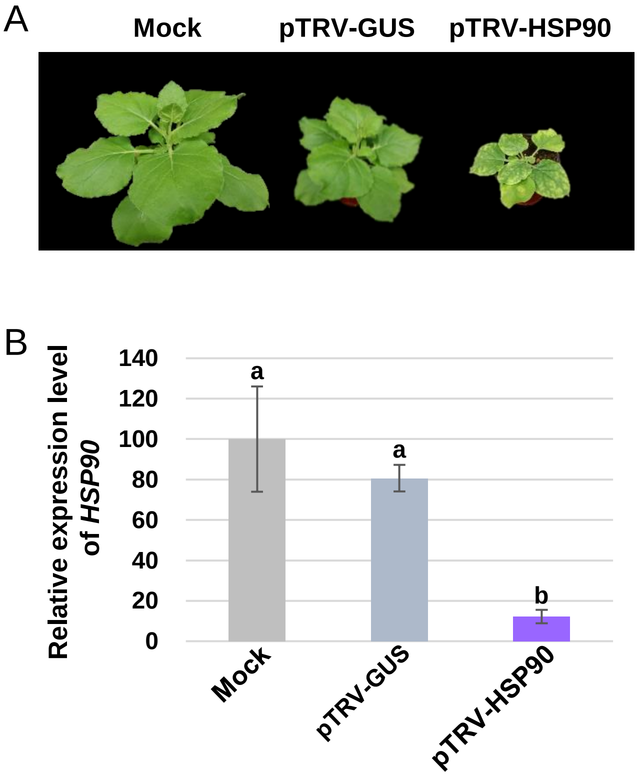


**Supplementary Fig. S1.** Phenotypes of *HSP90*-silenced *N. benthamiana* plants. Two-week-old *N. benthamiana* plants were agroinfiltrated with mock (*Agrobacterium* containing no binary vector), pTRV-GUS, or pTRV-GUS. The plants inoculated with mock or pTRV-GUS served as negative controls. *HSP90* silencing caused strong growth inhibition and leaf yellowing in *N. benthamiana* plants. The plants were photographed at 21 dpi with the TRV-based VISG constructs. (B) RT-qPCR analysis of *HSP90* expression. Total RNA isolated from the systemic leaves of *N. benthamiana* plants at 21 dpi was analyzed using RT-qPCR to assess the silencing efficiency of *HSP90*.

### Supplementary Figure S2

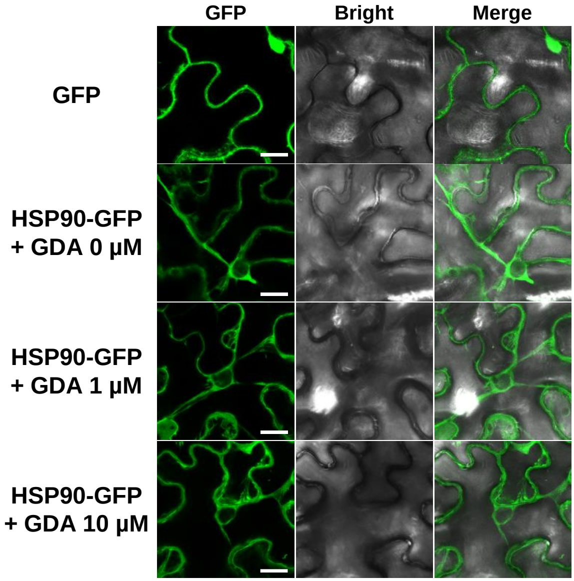


**Supplementary Fig. S2.** Effects of inhibiting the chaperone function of HSP90 on its subcellular localization. HSP90-GFP was expressed in *N. benthamiana* leaves using an *Agrobacterium*-mediated gene expression method. After 36 h, 0, 1, or 10 μM GDA was syringe-infiltrated into the agroinfiltrated leaf area. The subcellular localization of HSP90-GFP in the epidermal cells was observed using confocal microscopy at 3 dpi. Bar = 20μm.

Supplementary Table S1. Primers used in this study.

| Primer | Primer sequence (5' to 3') | Purpose |
|---|---|---|
| VP37-EcoRI-Fw | ACGAATTCATGAATGAGGCAAATATCAC | To construct pGBKT7-VP37 |
| VP53/37-BamHI-Stop-Rv | CGGGATCCTATTGACCATATCTATAATC | |
| VP53-EcoRI-Fw | ACGAATTCATGCGTCCCGAACTTGTTG | To construct pGBKT7-VP53 |
| VP53/37-BamHI-Stop-Rv | ACAAGTTCGGGACGTTATACAGCATGTTCA | |
| HSP90-Fw2 | GATGGCGGAGGCGGAGACGTT | To construct pGADT7-HSP90 |
| HSP90-Rv2 | CCGCTCGAGTTAGTCAACTTCCTCCATCTTGCT | |
| HSP90-T7-Fw | TTGAGATCTTTAATACGACTCACTATA | To amplify HSP90-Flag templates for *in vitro* coupled transcription/translation assays |
| 3E-PolyA-Rv | TTTTTTTTTTTTTTTTTTTTTTTTTTTTTTTTGGGTTATGCTA GTTATGCGG | |
| VP37-T7-Fw | TTGGAATTTGTAATACGACTCACTATA | To amplify VP37-Myc templates for *in vitro* coupled transcription/translation assays |
| 3E-PolyA-Rv | TTTTTTTTTTTTTTTTTTTTTTTTTTTTTTTTGGGTTATGCTA GTTATGCGG | |
| VP37-SalI-1-Fw | ACGCGTCGACATGAATGAGGCAAATATCAC | To construct pENTR$^{TM}$1A-VP37 |
| VP37-XhoI-1014-Rv | CCGCTCGAGGGTTATTGACCATATCTATAATC | |
| HSP90-SalI-1-Fw | ACGCGTCGACATGGCGGAGGCGGAGACG | To construct pENTR$^{TM}$1A-HSP90 |
| HSP90-Rv3 | ACTCGCTCGAGTTARTCAACTTCCTCCATCTTGCT | |
| HSP90-Fw1 | ATGGCGGAGGCAGAGACGTT | To construct PZP-HSP90-GFP |
| HSP90-SpeI-Rv | AGGACTAGTTTAGTCAACTTCCTCCATCTTG | |
| GUS-1-BglII-Fw | GAAGATCTATGTTACGTCCTGTAGAAACCCCA | To construct pBBWV2-R2-GUS |
| GUS-1809-AvrII-Rv | GATCCTAGGTTGTTTGCCTCCCTGCTGCGG | |
| HSP90-Vigs-SalI-223-Fw | TACGCGTCGACACCCTGACTATTATTGACAGTGG | To construct pTRV2-HSP90 |
| HSP90-Vigs-SalI-523-Rv | TACGCGTCGACTAATTTTTGTACCCCTGCCAAGGTTC | |
